# Bayesball: Bayesian Integration in Professional Baseball Batters

**DOI:** 10.1101/2022.10.12.511934

**Authors:** Justin A. Brantley, Konrad P. Körding

## Abstract

Pitchers in baseball throw the ball with such high velocity and varying movement that batters only have a few hundred milliseconds to estimate whether to swing and how high to swing. Slight deviations in the contact point on the ball can result in weakly hit balls that do not result in opportunities for batters to score. Even before the pitcher releases the ball, the batter has some belief (an estimated distribution–a ‘prior’), of where the ball may land in the strike zone. Batters will update this prior belief with information from observing the pitch (the ‘likelihood’) to calculate their final estimate (the ‘posterior’). These models of behavior, called Bayesian models within movement science, predict that players will estimate a final ball position by combining prior information with observation of the pitch in a way that weights each information source relative to the uncertainty. Here we test this model using data from more than a million samples from professional baseball. Moreover, as predicted by a Bayesian model, we show that a batter’s estimate of where to swing is biased towards the prior when prior information is available (‘pitch tipping’), and biased towards the likelihood in the case of pitches with high movement uncertainty. These results demonstrate that Bayesian ideas are relevant well beyond laboratory experiments and matter for real-world movements.

## Introduction

*You can’t think and hit the ball at the same time*

—**Yogi Berra**

It is remarkable that baseball batters can successfully hit the ball so often despite the uncertainty in pitched balls. Professional pitchers throw the ball at velocities up to 105 mph^1^ over a distance of 60.5 feet (Figure 1A), leaving batters approximately 400 milliseconds to perceive the ball, decide on an action, and swing^2^. With such limited perceptual systems, batters can only observe the ball for a short period of time until the brain needs to make a decision. It takes approximately 100-ms to observe the ball and an additional 150-ms to physically swing the bat^2^, leaving 150–250-ms for the batter to decide whether or not to swing. When deciding to swing, batters must account for uncertainty about the specific trajectory of the pitched ball. A large portion of this uncertainty is due to the pitcher varying the arm angle from which the ball is the thrown, the velocity, and by spinning the ball in various ways to introduce curved trajectories that are hard to predict (i.e., different pitch types). Pitchers will often try to further increase the challenge for the batter by ‘tunneling’ their pitches—a technique where they throw sequential pitches with varying velocity and spin profiles in a way that their trajectories are nearly identical early in their flight, but diverge sharply near the plate^3–5^. Finally, a pitcher’s throwing mechanics, their height, arm length, and stride during the throw can all impact the ball’s ‘perceived’ velocity, where balls released later in the delivery and closer to the plate may appear faster than the same pitch from a different pitcher with shorter body proportions^6,7^. Despite differences in movements between pitchers, individual pitchers try to make distinguishing between these trajectories difficult by moving their bodies in similar ways despite introducing multiple sources of uncertainty (e.g., velocity, spin, location thrown—Figure 1A-B).

**Figure 1.**
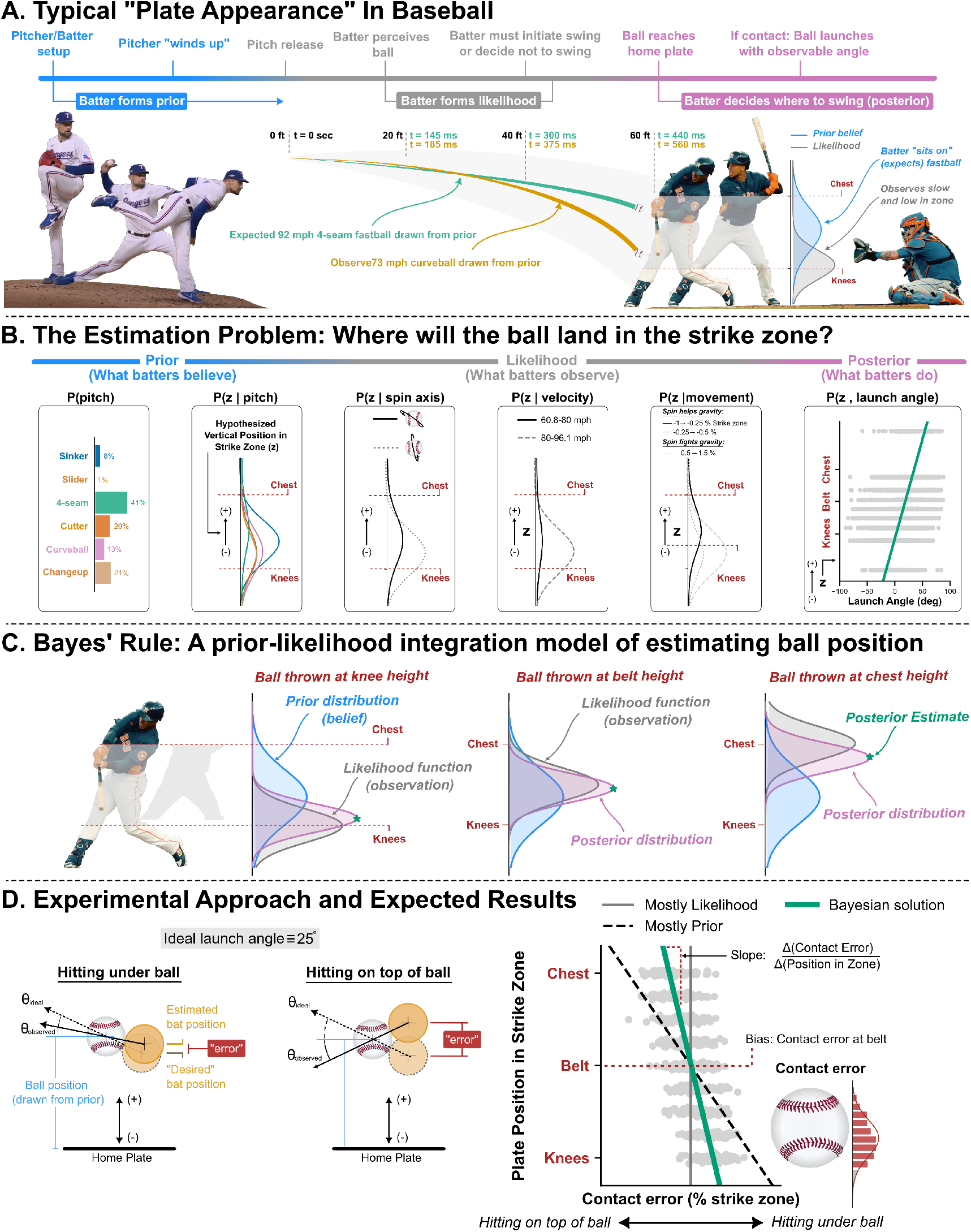
Testing Bayesian ideas by analyzing Baseball data. **A**. Lead up to a typical plate appearance, a batter may form a prior over where the ball will land in the strike zone. They will subsequently observe the ball’s trajectory and form a likelihood distribution for the final location, which represents the probability given the observation of the pitch. **B**. A batter’s belief about where the ball will land is based on the pitcher’s unique pitch behavior, such as pitch selection, velocity, spin rate, and movement. **C**. Bayes’ rule allows for the combination of prior knowledge with observations to make an estimate, called the posterior. **D**. *Left:* We test whether batters behave in a Bayesian way by estimating the contact error on batted balls (hitting under/on top of ball). *Right:* Our theoretical model (simulated data) illustrates a Bayesian solution (green line), a mostly likelihood model (solid line), and a mostly prior model (dashed line).

At the moment when the ball leaves the pitcher’s hand, the batter can at best estimate a probability distribution of where the ball may end up. This distribution is called the prior (Figure 1A, right, shown in blue) and is one of the pieces of the problem of estimating where the ball will land. The second part of the problem is extracting information about where the ball will be thrown based on observing the trajectory: the brain could form a distribution (Figure 1A, right, shown in grey) representing how probable the observations of the pitched balls are under which assumption of the thrown ball trajectory (ignoring the prior). While this information is noisy due to perturbations on the thrown ball, information from seeing the ball is clearly useful and may be summarized by the so-called likelihood function (Figure 1A, right, shown in grey). This defines how compatible each observed ball trajectory is with a potential height within the strike zone. Despite all of these uncertainties, batters usually make reasonable estimations of the ball’s trajectory and are still able to achieve hits approximately 25% of the time on average across all batters, with some exceptional players reaching rates of over 30%.

How can a batter predict a ball’s trajectory so well that they contact the ball successfully? More generally, we may ask how baseball batters estimate the pitched ball’s trajectory if all they have is noisy prior information and the noisy sensory information from watching the pitched ball? Several hypotheses may explain how batters deal with pitch uncertainty during batting. The first is by only considering prior information. By generating a distribution over pitches, the batter might then select some action (prior to the pitcher throwing the ball) that maximizes the probability of contact under this distribution. Importantly, this hypothesis assumes that the batter makes no adjustments based on the observations of the thrown pitch. A second approach is that they could rely exclusively on observable data from each pitch without using any prior knowledge for their batting estimate; however, this hypothesis is unlikely to accurately describe a their behavior since it assumes that batters do not make estimates about the visual uncertainty based on prior information. A third strategy, which follows Bayesian statistics, predicts that a batter will combine prior information with the observation to make a prediction about the oncoming pitch. In the language of Bayesian statistics, we might say that a batter combines their prior beliefs about a pitch, **P**(*A*), with the likelihood, or probability of the observed pitch given that the belief is true, **P**(*B*|*A*). They then make a prediction, **P**(*A*|*B*), which is the posterior probability of the pitch after taking into account the observation of the pitch (Figure 1C). Formally, we express this as

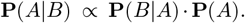

The result is a situation where what we know from observing the pitcher (prior) and what we observed during the ball’s flight (likelihood) are combined based on the level of uncertainty in each. For example, when vision of the ball is very good (e.g. because the ball is slow or the spin pattern is clear), batters should rely mostly on how they see the ball fly. On the other hand, when the pitcher is more predictable, batters should rely more on their prior knowledge. This approach is favorable in that it allows for the use of prior information about pitch behavior while permitting flexibility in the estimate by accounting for uncertainty in both the prior and observation. Practically speaking, this is expected as we know that batters study opposing pitchers in advance of the game (prior information) and that batters often achieve optimal outcomes despite the varying sources of uncertainty on the pitch. To test Bayesian ideas we need to know about

1. the existence of the prior distribution (e.g. from the distribution of heights of pitches within the strike zone);
2. the results of a perturbation (the final vertical position of the ball); and, importantly,
3. we need to know the error made for each estimate.

The first two can be directly measured, however, we must make an approximation for the estimation error on batted balls.

A Bayesian approach to sensorimotor control has been extensively documented through decades of laboratory studies on movement and decision making^8–13^. A common experimental approach to test this hypothesis for motor control is a 2-dimensional reaching task, e.g., using a robotic manipulandum, a tracking system, or a computer cursor to capture the movement behavior, where the subjects are asked to reach towards a target while a perturbation is applied to the movement path. The perturbations are drawn from some prior distribution and feedback with varying levels of uncertainty is provided to the subjects about their action. Since the correct path can only be noisily observed (through the feedback with varying uncertainty), the subject should theoretically alter their movement path to correctly execute the task based only on their prior information and the noisy feedback. Indeed, subjects behave in a way consistent with Bayes’ rule, where they rely more on prior knowledge when observing feedback with high uncertainty and rely more on the feedback when it contains lower uncertainty. While these studies suggest that the brain implements some form of Bayesian integration for managing uncertainty, “Bayesian brain” behavior has scarcely been documented in real world data. Moreover in addition to observing this behavior, we ask, does Bayesian behavior matter for real world movements? Here we use the batting behavior of professional baseball players to test if humans use Bayesian approaches in the real world. Specifically, we hypothesize that batters combine prior information about the pitch with observations during each time they bat to achieve contact, and that this behavior can be explained using a Bayesian framework (simulated example in Figure 1D, right). We find that the behavior observed in the baseball data is consistent with laboratory experiments and implies that batters combine prior information with noisy observations to manage pitch uncertainty in a way that is consistent with Bayesian statistics. Furthermore, when batters have prior information about the oncoming pitch, our results show that they rely more on the prior knowledge. Moreover, we show that for certain pitch types with high probability yet high behavioral uncertainty (that is, batters know what pitch is coming but its movement is noisy), batters rely nearly completely on observations about the oncoming pitch while ignoring the uninformative prior knowledge associated with the pitch type. These results demonstrate Bayesian behavior in a real world skilled movement task.

## Results

We test if baseball batters employ a strategy for managing pitch uncertainty that can be explained by Bayesian statistics. We gathered data from MLB’s publicly accessible data clearinghouse, Statcast^14^, for MLB regular and postseason games from 2015-2023 (5,882,670 pitches before processing, 1,686,238 pitches retained for analysis). Within this data set we can distinguish different pitch types, giving us variables that may affect the prior uncertainty of pitches. With this, we can thus approach asking questions about uncertainty processing by professional batters.

Estimating vertical pitch position is challenging for the batter because it varies from pitch to pitch. We directly have this information in our data since it is measured for each thrown pitch. The overall distribution of the vertical pitch position was roughly normally distributed, *N* (*µ* = 0.46, *σ* = 0.33), with approximately 87% located within the vertical limits of the strike zone (Figure 2A). This distribution of vertical pitch position is crucial as it defines the distribution of relevant variables to be estimated. Pitchers use the entire strike zone, and even beyond, to make the estimation task difficult for the batter.

**Figure 2.**
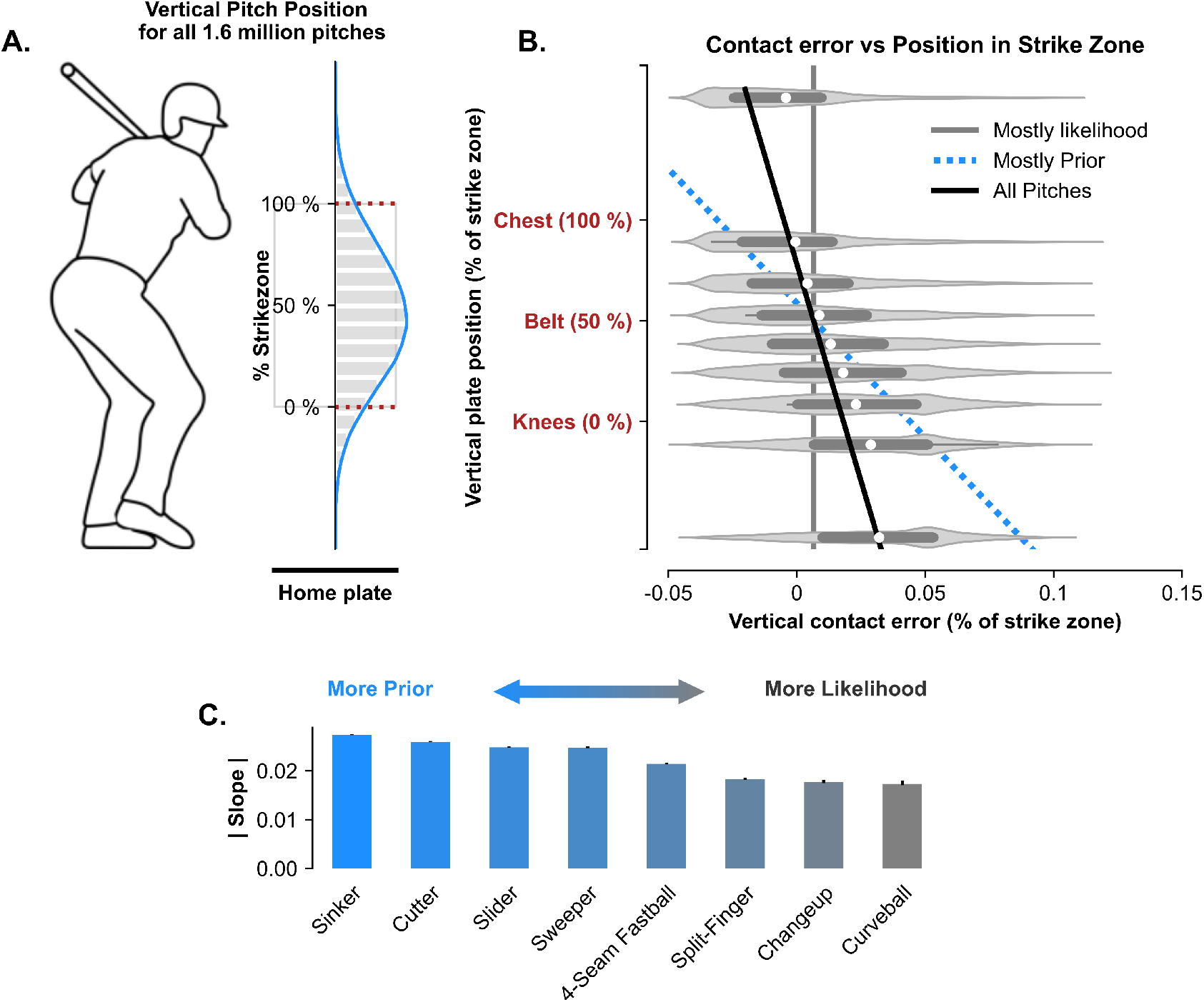
batters combine prior information with noisy observations to manage pitch uncertainty. **A**. Distribution of vertical pitch location within the strike zone for all 1 million pitches. **B**. Vertical contact error as a function of vertical plate position. The solid gray line indicates a mostly likelihood solution and the dotted line indicates a mostly prior solution. The blue line indicates a Bayes-like solution across all pitch types. **C**. The slope of each pitch type. The terms sinker, cutter, slider, sweeper, 4-seam fastball, split-finger, changeup, and curveball are all specific pitch types with varying speed and movement profiles that are likely to impact how batter’s employ a prior-likelihood integration strategy. **Note:** The reported slope is for the regression model when the vertical plate position is treated as the explanatory variable and the contact error is the response. The axes are switched for visual clarity.

The basic prediction of Bayesian statistics is that the further up the ball is within the strike zone, the more strongly the batter should estimate a position biased downward and vice versa. We can test this idea by plotting the inferred error as a function of the position within the strike zone. We computed the estimate of the error as the difference between the “ideal” contact point on the ball (i.e., the point on the ball that results in a launch angle that maximizes the probability of a home run) and the observed contact point (example shown in Figure 1D). Indeed, we find a systematic correlation between position and error as predicted by the theory (Figure 2B), suggesting a Bayesian strategy for integration where both prior and likelihood are used for estimation. Additionally, we find that the level of reliance on each varies by pitch type.

### There is no evidence of learning effects

It is commonly thought that batters are able to learn over the course of a single plate appearance after seeing several pitches in a row. We used our model to look at consecutive pitches within the same plate appearance (i.e., same pitcher, batter, game, and inning) in which contact was made on each of those sequential pitches. We hypothesized that if batters learn from consecutive pitches, our model would reveal a systematic relationship between the contact error at time, *t* (current pitch), and the vertical plate position of prior pitches. We find that the location in the strike zone on the two previous pitches has no impact on the error for the current pitch (Figure S1), implying that batters are unable or choose not to learn from consecutive pitches. In addition to learning, we observe that batters are slightly correlated in the error on consecutive batted balls (Figure S2). This effect is very weak and is likely attributable to internal error processes, such as motor error or changes in strategy. These results make sense for professional baseball; if batters tried to learn from consecutive pitches, pitchers could easily exploit the known temporal dependency in their batting strategy.

### External cues impact the strength of the prior

Most major league pitchers throw an average of 4-5 pitches, usually with varying movement patterns and velocities to maximize uncertainty when sequencing pitch types. However, pitchers usually rely on some pitches more than others (particularly depending on the handedness of the batter). Information about a pitcher’s pitch usage (and related pitch behavior) is readily available to teams and players, who presumably can then use this information to generate some form of a prior for each pitcher. However, there are cases in which external information can be relayed to the batters about oncoming pitches (e.g., pitch tipping, decoding of signs, and cheating), thus influencing the prior. In this study, we consider a specific case in which the decoding scheme is known.

In a game on October 10, 2019^*^, Tyler Glasnow, an elite pitcher for the Tampa Bay Rays, was purportedly tipping his pitches based on how he positioned his glove along the length of his torso. In 2019, Tyler Glasnow threw three pitch types: the 4-seam fastball (67%, 96.9 *±* 0.4 mph), a curveball (29.3%, 83.5 *±* 0.2 mph), and a changeup (3.5%, 92.9 *±* 0.9 mph); thus, there is a relatively strong prior indicating that two pitches are more probable. However, due to the significantly varying behavior of the 4-seam fastball and curveball, batters struggled to hit well against his pitches. In this particular game, it became quickly apparent that batters were able to predict which pitch was being thrown simply based on the position of the glove along his torso (Figure 3B). When the glove was high (in line with his neck), he was planning to throw a 4-seam fastball, whereas when the glove was lower (around the logo on his jersey) he was planning to throw a curveball. When comparing the data from this particular game to all other games throughout the 2019 season, batters relied significantly more on the prior when making predictions as indicated by the slope in Figure 3B (right). Thus, these data suggest that external information (pitch tipping) influences the weight that batters place on the prior and likelihood when making inferences. In addition to the above example, we have included a second example of pitch tipping in the supplementary materials to further strengthen our claim that pitch tipping biases batters toward the prior (Figure S3).

**Figure 3.**
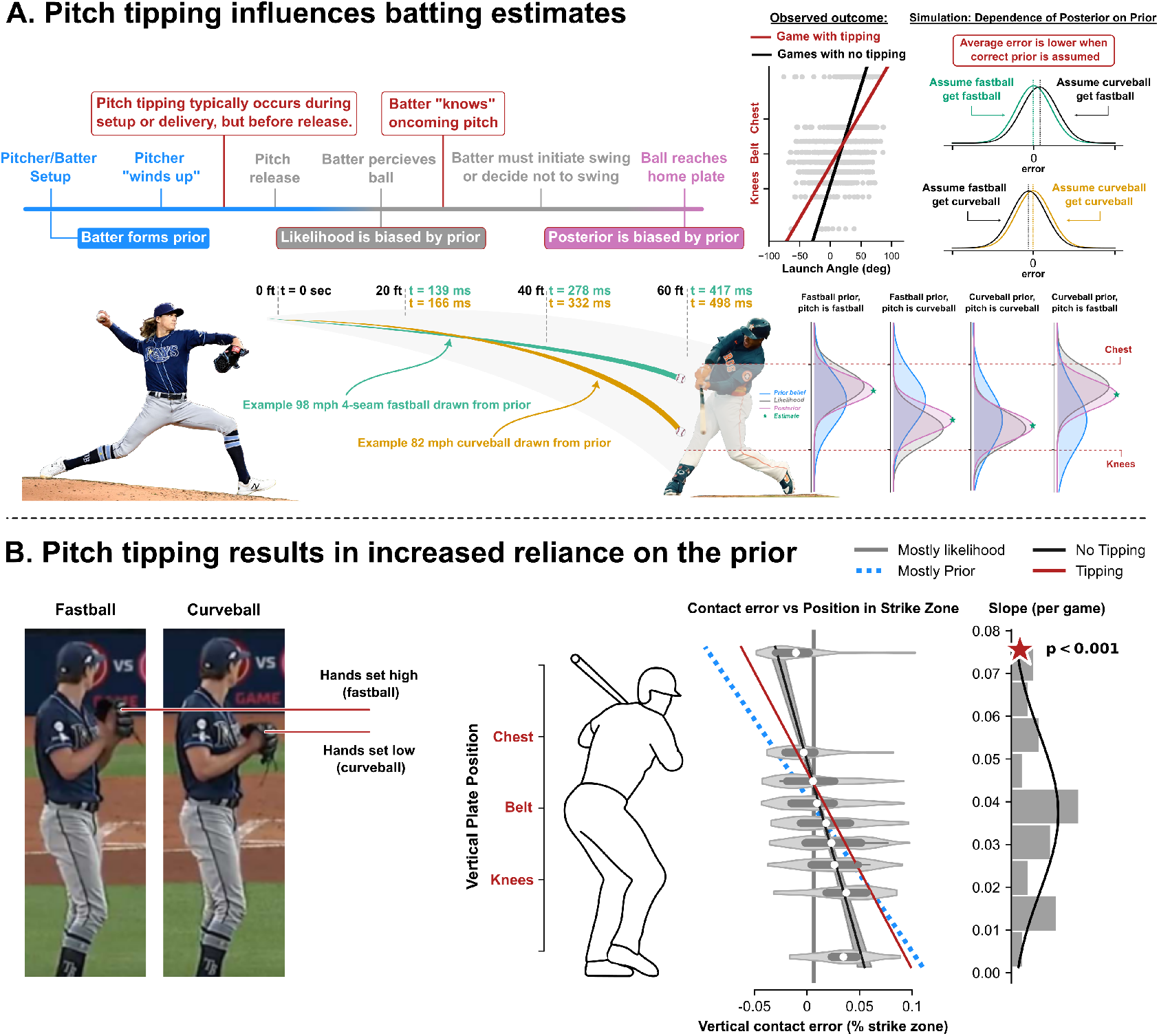
Pitch tipping results in a batting estimate biased toward the prior. **A**. In the timeline of a single plate appearance, pitch tipping occurs before ball release, either during the setup or during the wind-up. This results in pitchers having prior knowledge about the oncoming pitch. The error between the posterior estimate and the actual ball position is lower when the correct prior is chosen (simulation). **B**. *Left:* In some cases, the posture of a pitcher may give away the planned pitch. Here, when the glove is high, the pitcher is throwing a 4-seam fastball and when it is lower, the oncoming pitch is a curveball. In this case, we should find more reliance on the now improved prior. *Right:* As predicted, we find that the slope is higher and thus that the batters rely more on the prior.

### Pitches with weak priors but high movement uncertainty facilitate dependence on the likelihood

Knuckleball pitch behavior is characterized by erratic and unpredictable movements that make it challenging to predict the ball trajectory due to high movement uncertainty (a phenomenon observed in many ball sports). In the years 2015-2021, there were a total of 3,054 knuckleballs thrown across 8 pitchers. Among these pitchers, two pitchers account for 97.6% of all knuckleballs thrown, with one pitcher throwing knuckleballs 83.3% of the time (R.A. Dickey^16^; 2492 total pitches) and the other throwing them 74.7% (Steven Wright^17^; 1210 total pitches). Thus, when either of these pitchers was throwing the ball, there was a strong prior for what pitch the batter was likely to see. However, due to the highly erratic behavior of the knuckleballs, the high movement uncertainty renders the prior very weak. As shown in Figure 4D, the knuckleball results in a slope that trends towards zero, indicating a high reliance on the likelihood, despite the potentially strong prior of knowing that the particular pitch type will be thrown. In laboratory settings, this pitch is analogous to a condition in which both the behavior prior and likelihood are drawn from a wide distribution, or a distribution with high uncertainty. In this case, the “weak” prior knowledge (due to high movement uncertainty) results in a posterior estimate having greater reliance on the more “reliable” observed information, despite also being drawn from a wide distribution. This can either be interpreted as the batter ignoring the prior altogether, or simply assuming a flat prior with uniform probability for any location within the strike zone.

**Figure 4.**
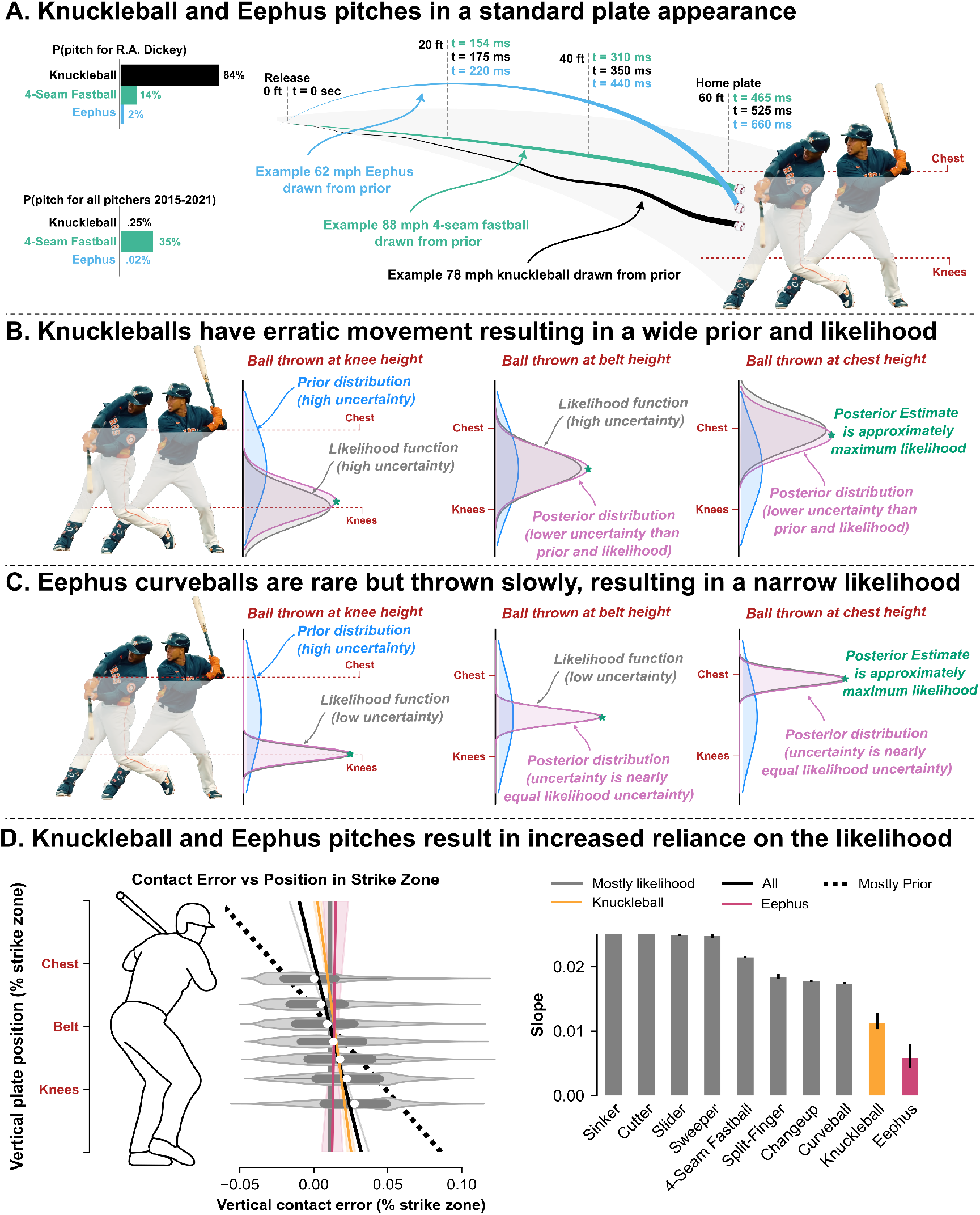
Pitches with weak priors facilitate dependence on the likelihood. **A**. The movement path of the knuckleball (shown in black) is highly erratic leading to high uncertainty. The eephus has a slow arcing movement profile (shown in light blue) and occurs very rarely in professional games. **B**. Knuckleballs have a noisy likelihood but a noisier prior, thus it cannot be predicted easily. The posterior estimate is theorized to be approximately the maximum likelihood solution. **C**. The Eephus is a slow pitch and thus easily observable, however, the prior is weak since it is an uncommon pitch. **D**. *Left:* Vertical contact error as a function of vertical plate position. The solid gray line indicates a mostly likelihood solution, the black dotted line indicates a mostly prior solution, and the black line is for all pitches. The orange and pink represent the knuckleball and eephus, respectively. *Right:* The slope and bias of each pitch type. **Note:** The reported slope is for the regression model when the vertical plate position is treated as the explanatory variable and the contact error is the response. The axes are switched for visual clarity.

### Pitches with weak priors but low movement uncertainty facilitate dependence on the likelihood

Another unusual, yet interesting baseball pitch is the ee-phus, which is a slow, often looping, curveball that is thrown at a much lower velocity than all other pitches. The eephus is a rare pitch, thus resulting in a weak prior for batters when predicting a pitch type (although some pitchers are known for throwing them at a higher rate, it is still uncommon). How then do batters make inferences on a pitch that is so rarely seen? Two distinct traits about the eephus make it an “easy” pitch to hit: the flight path of an eephus curveball tends to follow an arc trajectory and the velocity of the eephus ranges from the low 50 mph to high 60 mph range–velocities otherwise commonly seen by youth players. In sensorimotor experiments, this pitch is analogous to the case of a wide prior and a narrow likelihood and results in subjects relying on the feedback, or likelihood. As expected, our results reveal a slope close to zero (Figure 4D), indicating almost complete reliance on the likelihood due to the weak prior and low observation uncertainty. In other words, the slow velocity and arcing movement allow for significantly more time to observe the pitch and thus a near complete reliance on the observation.

## Discussion

Here we asked if professional baseball batters use a Bayesian approach when hitting a pitched baseball. We used over 1 million pitches from Major League Baseball games to estimate the error on batted balls and their dependence on the pitch. We found a systematic dependence of contact errors on pitch position within the zone, with each pitch type being accompanied by its own level of uncertainty. Moreover, we found that this is strongly interacting with pitch tipping, where pitcher behavior gives batters better priors. We also observed that when facing highly uncertain or uncommon pitches, the priors are largely ignored resulting in a posterior estimate approximately at the maximum likelihood estimate. These effects are well predicted by a conceptual Bayesian prior-likelihood integration model. Our results suggest that batters rely on prior knowledge and real-time observations (likelihood) when making predictions for batting. In laboratory studies, the observed error, or deviation, can be related back to the true shift imposed by experimenters. When the batter sees the pitcher prior to the throw, they can form an expectation of where the ball will land, e.g. by looking at the pitcher, thus drawing from a distribution known as the prior in Bayesian terms. However, this prior will be different from trial to trial and it cannot be measured directly. The batter’s estimate of where the ball will land can then be seen as a way of improving the estimate based on seeing the ball’s trajectory. However, we can not measure this prior distribution since it largely depends on what is going on in the batter’s brain. But, any ball’s position can be seen as traveling towards the mean of the prior and then having an error relative to that. This error is what Bayesian statistics operates upon. As such, we cannot predict the exact dependence of batting errors on the pitched ball trajectories. However, what we can predict is the trend coming from Bayesian statistics: a ball landing lower should lead to an error further up on the ball. This dependence should be represented by a steeper slope if the prior information is better and shallower if the likelihood information is better. For our analysis we have to approximate the calculation of batting contact error, which we consider to be the deviation from the “true shift”. Our analysis uses a simple 1D geometrical model that estimates the deviation in the vertical axis based on the launch angle off of the bat. This model does not account for the tilt or fan of the bat, which will have an impact on the contact point on the ball. Additionally, we assume that the attack angle of the bat and the center-line angle of the ball are aligned such that the contact is purely normal to the bat surface. Due to the limitations of our data, we are not able to explicitly address these assumptions in our model; however, we believe that this model is a fair first-order approximation of the bat-ball contact and that the behavior observed in our data is persistent across the millions of pitches incorporated in our analysis. There is no reason to believe that the relevant angles would systematically depend on factors expected to influence the prior. As tracking technology continues to advance, future research can use a full calculation based on precise tracking of the ball and bat. One important question to consider is whether biomechanics alone can explain the results of our analysis. Could it be that our finding that error is related to height within the zone is simply explained by the swing starting at a point where the batter does not have enough time to raise or lower the bat sufficiently? That interpretation seems unlikely to us based on both existing knowledge and our data. It is well established that batters are still able to achieve hits, including home runs, when facing extremely high velocity pitches, even when the ball is high or low. At these fast pitch velocities, the necessary bat velocity required to hit the ball is far too high if the batter waits until the pitch has been fully perceived. In order to hit the ball, the batter must initiate the swing prior to recognizing the ball by relying on some prior belief about the pitch. Moreover, the assumption of the effect being entirely biomechanical can neither explain why tipping leads to what appears to be stronger priors. The effect of a more localized prior is similar to the effect of a batter being weaker, in that both effects predict that larger deviations of ball positions are associated with larger errors. However, the explanation in terms of priors naturally predicts that a) tipping produces a stronger prior and thus larger deviations, and b) that knuckleballs are weakly represented in the prior and thus smaller deviations. The biomechanical explanation seems unlikely: Why would batters be weaker in the case of tipping and stronger for knuckleballs? As such, the most parsimonious explanation of our data remains the Bayesian one. We do not know precisely what batters are actually trying to do. For example, batters behave differently based on external factors, such as the pitch count, the score, the number of outs, or based on directions provided by the coaches (e.g., “do not swing at the first pitch”). Nonetheless, we believe that these external considerations do not strongly impact the prior or the batter’s behavior of integrating the prior and likelihood in a Bayesian manner. Instead, this is likely to only impact the batters interpretation of “optimal” for that specific scenario. For example, a batter attempting to either simply get on base (e.g., hit a single) or to hit a sacrifice fly to score a runner both result in optimal launch angles not equal to the optimal launch angle for hitting a home run. While situational batting is important, we do not see how it could explain our data. Explaining baseball behavior in terms of probabilistic ideas has received some attention in the sports science literature. The idea that ball trajectory estimation requires priors and likelihoods has been discussed as parts of the process of batting^18–21^ and sports more generally^22^. Our results empirically demonstrate that batters rely on both prior information and observable information when hitting and that the reliability of these information sources can be weighed by their relative uncertainty. Laboratory studies have also shown that the various sources of information that we discuss empirically matter to athletes^18–20,23–28^. The contribution of our study was testing this set of ideas directly on a large database of professional batting in major league baseball games. In the space of sensorimotor integration and motor control, it has been well observed that subjects use priors and noisy feedback based on their respective uncertainties in a way that is consistent with Bayesian statistics. This type of experimental approach, which is typically conducted by introducing some type of visuomotor or dynamic perturbation to a simple movement task, has been iterated in a number of environments, such as in force estimation, cue combination, motor adaptation, and a coin splash game in children. We were inspired by these studies to study baseball to see if this behavior can be observed in the real world. The location where the pitch lands in the strike zone in a way fulfills a similar role to the perturbation in the lab based studies. Just like in lab based studies, there is a prior distribution, with the slight difference that it is not entirely observable for us. In addition, there are factors that influence priors, for example, the case of knuckle-balls (relatively flat prior) and tipped pitches (relatively narrow prior) in our study. Moreover, the ball’s angle implicitly reveals the estimated position, in the same way as movement patterns reveal implied estimates in lab experiments. With this design we were able to find evidence of Bayesian behavior in professional sports. Our study brings Bayesian movement science into the real world. It opens the path towards computational motor control with big data from other professional sports, such as tennis^29^, penalty-kicks in soccer^30^, or squash^31^. The emergence of the technology of video based pose-tracking could help to strengthen and enable such analyses. More generally, these findings allow us to observe movements with high quality in the real world. This promises to allow us to bring lab based ideas with their conceptual sophistication to the real world and, ultimately, impart movement science with high ecological validity. The field of motor control needs to bring model based thinking to the real world, and maybe, Bayesian movement control, a large and active field largely confined to laboratories today, will improve athletic performance.

## Methods

### Data Acquisition

Our analysis was conducted using Python 3.10. All code used for data acquisition and analysis have been made openly available. We used the *pybaseball* library (https://github.com/jldbc/pybaseball) to obtain publicly accessible data from MLB’s data clearinghouse^14^. The on-field data are obtained through an acquisition tool known as Statcast^32^, which employs high-accuracy, high-speed cameras and radar to track the ball within the field of play, including pitch release velocity, pitch spin rate, ball position within the strike zone, and launch angle (among 91 features returned for each pitch^33^). Our initial query returned a total 5,882,670 pitches for the 2015-2023 seasons. We proceeded to exclude games from Spring training or exhibition games since these games are often used for players and teams to experiment with new strategies (e.g., a pitcher exploring a new pitch type, players at new positions, altered mechanics of the swing) and for teams to re-acclimate to playing after the off-season. We then removed pitches in which no contact was made between the bat and ball, including no swing or where the batter would swing and miss. Our resulting dataset included 1,686,238 pitches that were retained for analysis.

### Estimation of Bayesian Behavior

For our analysis, we used the final vertical position within the strike zone as the representation of the “true shift”, or the perturbation drawn from the prior distribution, which we computed by normalizing the true vertical height of the ball (measured in feet) by the height of the strike zone for the batter (computed per pitch). In a laboratory experiment, we would draw the “true shift” from a known prior distribution and then directly measure the deviation from that value based on the error in the batter’s estimate. However, since we do not have data that directly track the bat, we infer the contact point based on the launch angle of the batted ball. There are several factors that impact the launch angle of the batted ball, namely the center line angle of the ball and the attack angle, fan, and tilt of the bat^34^. Due to our inability to track the bat, we therefore assume that contact is purely normal such that there is no fan or tilt and the attack angle and the center line angle of the ball are aligned at contact. We assume a simple geometrical model, where the contact error, **e**_contact_, is computed as:

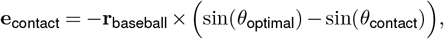

where **r**_baseball_ = 1.45-inches, *θ*_optimal_ = 25^*Ζ*^, and *θ*_contact_ is the launch angle observed when the bat contacts the ball. We then performed a final cleaning by removing extreme outliers in the data resulting from tracking errors. The data were binned into by discretizing the vertical position within the strike zone into 9 bins. We fit an ordinary least squares (OLS) model to fit the error on the batted ball as a function of the position within the strike zone for the aggregate pitch data (all pitch types) as well as for each unique pitch type (minimum number of pitches = 10,000). According to Bayesian statistics, we assume that a slope closer to zero indicates more reliance on the likelihood and an increasing slope indicates more reliance on the prior (Figure 1D). We performed this same analysis for the cases of knuckleballs and eephus pitches for the available number of pitches within the dataset.

### Pitch Tipping

For the case of pitch tipping we specifically identified pitches from the game during which the tipping occurred (November 10, 2019) as well as all other pitches thrown by Tyler Glasnow during the 2018-2020 seasons. We followed the same procedure as before by fitting an OLS model for each of the two conditions and used a t-test to determine the statistical significance of the slope during tipping compared to all other pitch appearances.

## Data/Code Availability

All code used for data acquisition, data analysis, and figure generating have been made available at https://github.com/KordingLab/Bayesball

## Acknowledgements

The authors would like to thank Dr. Brett Mensh for the significant editorial contributions. The authors would like to thank Professor Alan M. Nathan for the helpful discussions regarding the physics of the bat/ball contact model. Thank you to members of the Körding Lab for the feedback and discussion throughout the project.

## Author contributions

JAB and KPK conceived of the project together; JAB analyzed the data and created the figures; JAB and KPK wrote the manuscript.

## Supplementary Information

### No evidence of learning - supplementary figures

**Figure S1.**
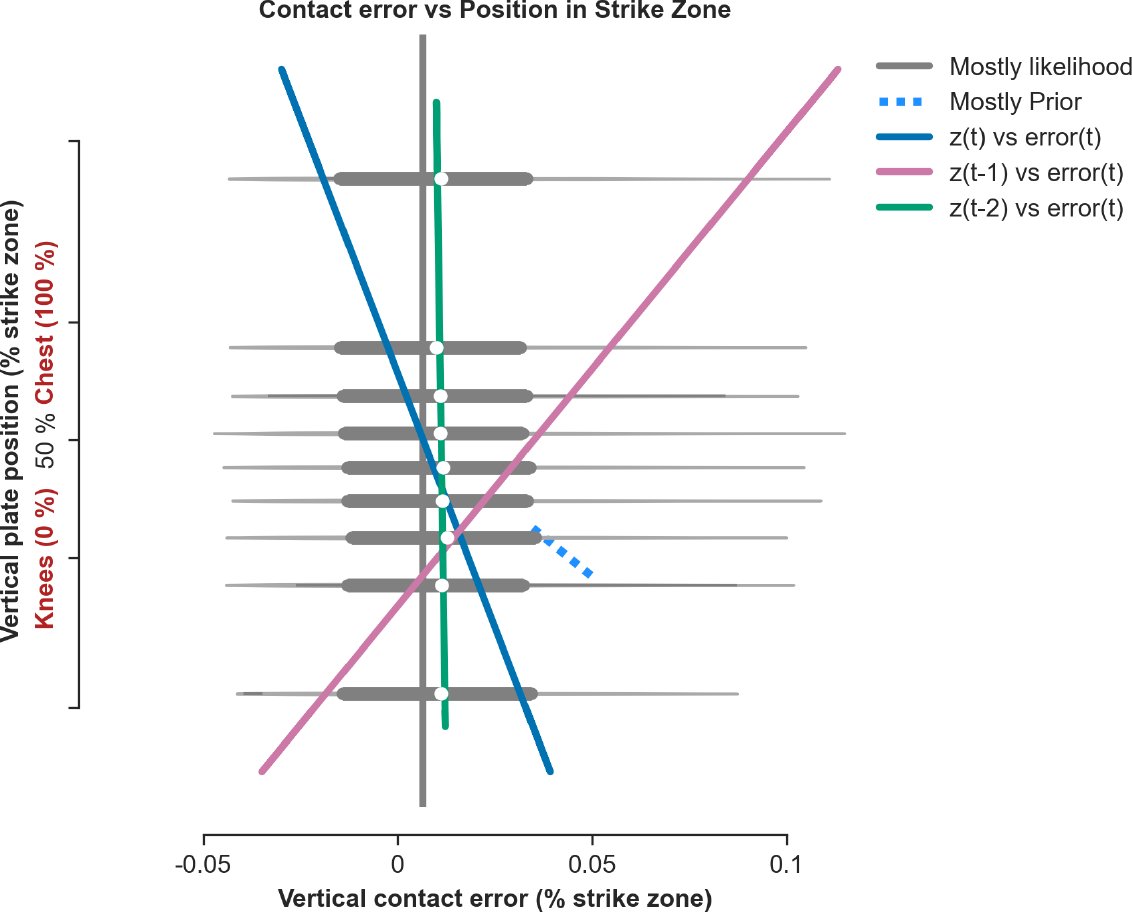
Vertical contact error as a function of vertical plate position at time t (current pitch), t-1 (previous pitch), and t-2 (two pitches previous). These results indicate that there is no evidence on learning over consecutive pitches.

**Figure S2.**
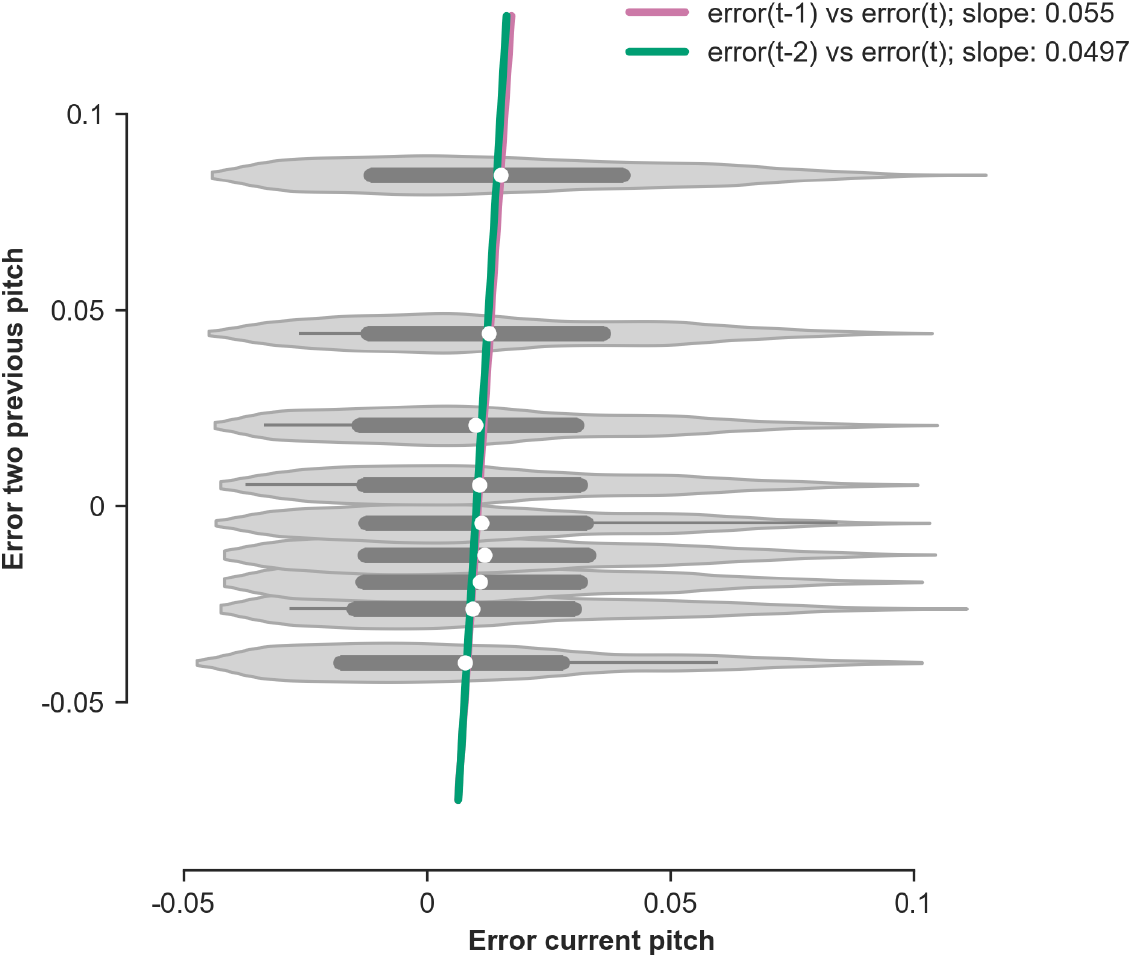
Error on batted ball at time t (current pitch) vs error at time t-1 (previous pitch) and t-2 (two pitches previous). These results suggest that there is a small amount of error between consecutive pitches that may arise from internal processes, such as motor error or changes in strategy.

### Additional example of pitch tipping

In our study, we consider the case of pitch tipping as an example of when batters are biased towards the prior, i.e., they know what is coming. To someone who is not a fan of the game, it can be challenging to believe that batters are capable to detecting such subtle queues by the pitcher that indicate what pitch might come next. While there are many documented examples in the past, we chose to highlight a second one that occurred within the time frame of our data. Additionally, the team who was batting (i.e., identifying the pitcher’s tips) is the same batting team as the example in the paper.

In a game on October 29, 2019 (game 6 of the World Series), Stephen Strasburg, an elite pitcher from the Washington Nationals, was the starting pitcher of the game. Similar to the first example, consecutive batters were achieving hits off of difficult to hit pitches. One of the key indicators of batters being able to identify pitches in advance is how well they time their swing and the quality of the contact (i.e., well struck balls). In this game, it was revealed that the pitchers hand movements in the glove were reveling what pitch he planned to throw. Importantly, this was an issue that he had had in the past and thus the coaches were able to identify the issue and help him better disguise his pitches for the remainder of the game. After the game, the pitcher acknowledged that he was indeed tipping his pitches, which further reinforces this idea that pitch tipping is a real and highly consequential phenomenon. In the following figure, we see that the same pattern of batters being biased toward the prior is observed for this case of tipping.

**Figure S3.**
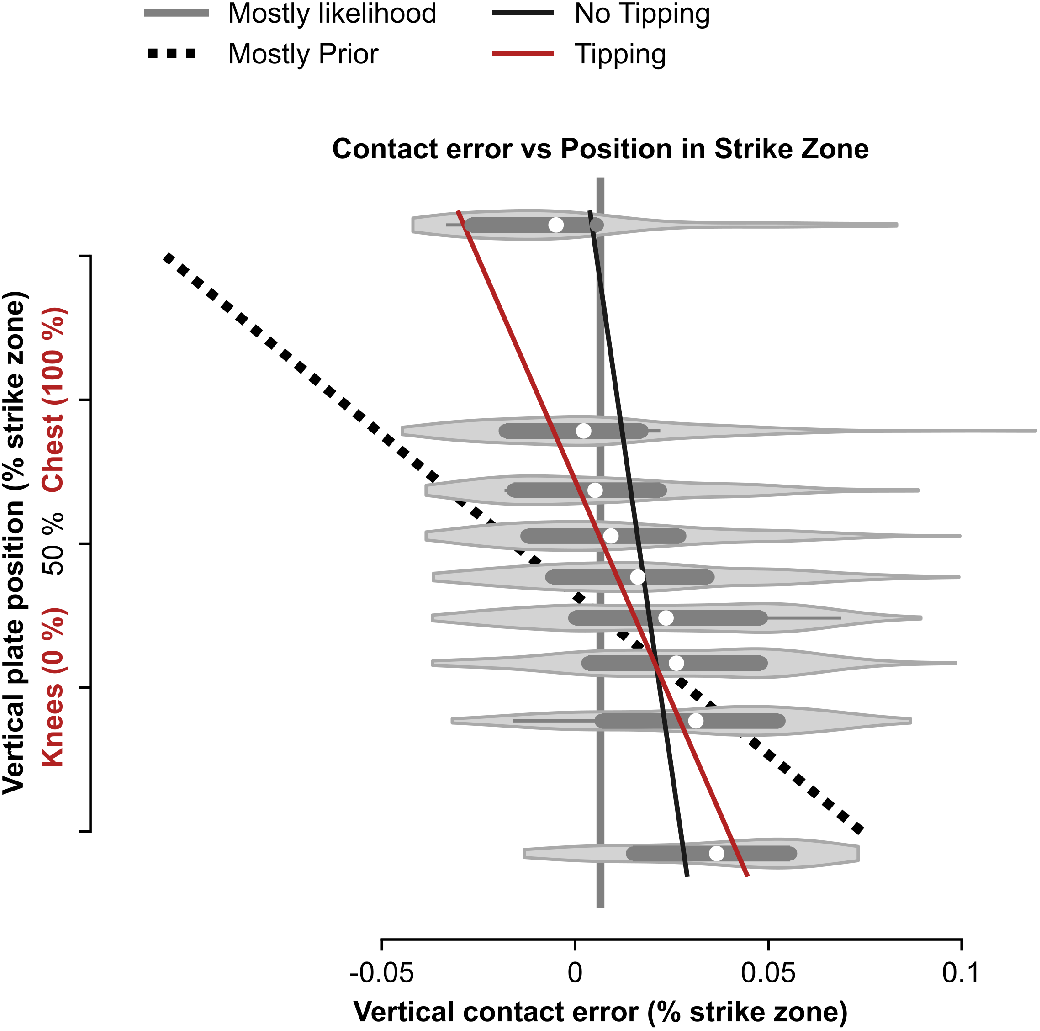
Pitch tipping results in a batting estimate biased toward the prior - Stephen Strasburg example.

This incident is further discussed in these articles and videos:

1. https://www.youtube.com/watch?v=aI52FQ23SW0
2. https://www.nbcsports.com/mlb/news/stephen-strasburg-says-he-was-tipping-pitches-in-first-inning-of-game-6
3. https://www.si.com/mlb/2019/10/30/stephen-strasburg-nationals-saved-season-world-series-game-6

## Tipping in other sports

The act of tipping can occur in many sports, such as,

- a baseball pitcher indicating which pitch he will throw,
- a soccer player indicating which direction they will kick the ball (e.g., during a free kick or penalty kick), or
- a tennis player indicating which way they will serve the ball.

For example, in 2017, the legendary tennis player, Andre Agassi, reveled that he had learned to defeat Boris Becker by watching his tongue. After studying Becker’s serve through film, he observed that Becker would unintentionally move his tongue in the direction that he planned to serve the ball. While we do not examine this example in our analysis, it serves as a strong example of when tipping occurs outside of the realm of baseball. This incident is well documented in the following resources:

- https://www.facebook.com/playerstribunefootball/videos/andre-agassi-how-i-beat-boris-becker/1249137535168463/
- https://www.businessinsider.com/andre-agassi-beat-boris-becker-watching-tongue-serves-2021-4

## Overview of Baseball

Baseball is a game played between two teams. The team playing in its home ballpark, or stadium, is called the home team, while the opposing team is the visiting, or away team. Baseball is played in turns, with each team alternating between offense and defense. The game takes place on a large field that resembles a triangle or diamond shape extending outward from a point known as home plate. Home plate is where batters stand to hit the ball.

There are three additional bases—first, second, and third—that form a diamond within the field. The area that includes the bases and extends inward toward home plate is known as the infield, while the area beyond the infield and within the two foul lines is known as the outfield. The main objective in baseball is for the offensive team (the batting team) to hit the ball and run around the bases to score runs. The defensive team tries to prevent this by getting the offensive team “out.” These terms will be discussed in more detail herein.

The game begins with the home team “on the field,” or playing defense. The team on the field is allowed nine players to defend against hit balls: the pitcher, the catcher, four infielders (first baseman, second baseman, shortstop, and third baseman), and three outfielders (left fielder, center fielder, and right fielder). The pitcher’s role is to throw the ball in a way that entices the batter to swing but makes it difficult for them to make good contact with the ball—or to hit it at all.

The batter’s job is to make contact with the ball as well as possible, aiming to hit it in a way that makes it challenging for the defensive players to stop it before the batter can reach base. If a batter strikes the ball with the optimal combination of launch angle and exit velocity, they increase the probability of hitting it over the back fence for a home run, which is one of the most desirable outcomes. Although not every batter is specifically aiming for a home run, batters generally strive to make hard contact. On average, a home run represents the best possible result of a well-hit ball.

The game is divided into innings, with each inning providing both teams a chance to hit (offense) and play in the field (defense). When the visiting team is hitting, it is referred to as the first half of the inning, or the top of the inning. When the teams switch, they enter the second half, or the bottom of the inning. The teams switch roles after the defensive team achieves three “outs.”

While this summary provides a short explanation for the game of baseball, the reader is encouraged to watch the following video for a more thorough visual explanation of the game: https://www.youtube.com/watch?v=tEckJtLgPIs

## Key Terms

**batter** Player on offense who hits the ball with the bat.

**estimation error** We define estimation error as the vertical error between the bat and the ‘optimal’ contact point on the ball to achieve a home run. The ‘optimal’ point is defined in the paper. The term ‘estimation error’ is not to be confused with an error in baseball, which specifically refers to a failed play that is deemed to be a misplay by the official scorer.

**hit** In baseball, a hit is a specific term that indicates that a batter made contact and was able to reach a base (or home run) without getting out. In this paper, when the bat and ball meet, we refer to this as ‘contact’ to distinguish from a formal ‘hit’ in baseball.

**inning** An inning is a subset of a full game of baseball that describes a one instance of both having a turn batting (offense) and playing in the field (defense; includes pitcher). A half-inning describes one of the teams taking a turn batting. A half-inning is over when the batting team gets three outs. A full inning is completed when each team has completed their half-inning. There are nine innings in a single game.

**pitch type** Pitchers vary the velocity, spin rate, and spin axis when they throw the ball to create curved trajectories. While behavior within a pitch type can vary from pitcher to pitcher, there certain velocity and movement that are characteristic of each pitch type.

Helpful videos summarizing pitch types: https://www.youtube.com/watch?v=0DFYJkneoMo https://www.youtube.com/watch?v=1FTFWzcgjHE

**4-seam fastball** A 4-seam fastball (or four-seamer) is the canonical ‘fastball’ in baseball. It is thrown in a way that minimizes movement and maximizes velocity, thus usally allowing pitchers to be more accurate with this pitch than most others. Four-seam fastballs usually have pure back spin, which can have the effect of fighting gravity over the flight path. A four-seam fastball thrown with very high rates of backspin can have the illusion of rising, making the pitch very difficult to hit.

**changeup** A changeup is a pitch that is designed to look like a fastball when gripped, but arrives to the plate with much slower velocity. The changeup is considered a deception pitch, where pitchers try to fool batters by throwing some combination of a fastball and changeup back to back.

**curveball** A curveball is a pitch that is usually thrown with signficant top spin, causing the ball to significantly drop in height as the ball reaches the batter. A well thrown curveball is often thrown following a fastball, where the curveball starts at with a similar flight path as a fastball, but arrives much slower and much lower in the zone

**cutter** A cutter is a variation of a fastball that is designed to have ‘glove side run’, meaning the ball will move left when thrown by a right handed pitcher and move right when thrown by a left handed pitcher

**eephus** An eephus, or eephus curveball, is a very rare pitch that is usually thrown in a high arcing path with low velocity. While few pitchers use an eephus regularly, several pitchers have used it as a tool in their pitch arsenal as a way to surprise batters

https://www.youtube.com/watch?v=Y7kJQrGKl-s

https://www.youtube.com/watch?v=ikLlRT2j7EQ

https://www.youtube.com/watch?v=xy5L4L5Q0nY

**knuckleball** A knuckleball is a pitch that is thrown in a way to minimize the spin to as near to zero spin as possible. This results in a flight path with erratic movements that makes the ball’s location unpredictable, making the ball difficult to hit

https://www.youtube.com/watch?v=qusYTWQFIF8

https://www.youtube.com/watch?v=35Sb5Jdtzz8

https://www.youtube.com/watch?v=e_Lhx2mfQ

**sinker** A sinker is a variation of a two-seam baseball that has late vertical movement or ‘arm side’ movement (moves right for right-handed pitcher and moves left for left-handed pitcher) or both movement profiles.

**slider** A slider has a similar ‘glove side’ run to a cutter, but is usually thrown with lower velocity and move vertical movement downwards. A slider is usually thrown with higher velocity and less vertical movement than a curveball.

**split-finger** A split-finger, sometimes called a splitter or split-finger fastball, typically has lower velocity than a four-seam fastball and significant downward movement late in the flight path. The splitter is wedged between two fingers on the throwing hand (creating a v-shape), which causes the ball to slide out of the hand with a lower velocity and low spin rate.

**sweeper** A sweeper, or sweeping slider, is a variation of a slider that has dramatic horizontal movement, usually moving 15” or more in the horizontal plane. The term, sweeper, is a relatively new term that was created to distinguish between a more traditional slider and the slider variation with huge horizontal movement. https://www.youtube.com/watch?v=z47hUrPKDG4

**pitched ball** In the context of this paper, we refer to any ball thrown by the pitcher to the batter as a pitched ball.

**pitcher** Defensive player who throws the ball to the batter. In the context of this paper, we refer to any ball thrown by the pitcher to the batter as a pitched ball.

**plate appearance** A plate appearance is any instance of a batter going up to bat, regardless of the outcome. This is in contrast to an ‘at-bat’, which is only counted in the score book when a plate appearance results in an out (including fielders choice), a hit, a strikeout, or fielding error

**statcast** Statcast is a camera and radar-based tracking system that is used for tracking players and the ball in many sports, including baseball and tennis. Statcast is used by MLB to track pitches (including velocity, spin rate, movement) and hitting (e.g., exit velocity, launch angle, batted ball distances). The data used in our paper are recorded using the Statcast system.

**strike zone** The space in front of the batter from approximately the knees to the sternum and the width of home plate.

## Footnotes

∗ Game 5, American League Division Series (ALDS)^15^

